# A geographic history of human genetic ancestry

**DOI:** 10.1101/2024.03.27.586858

**Authors:** Michael C. Grundler, Jonathan Terhorst, Gideon S. Bradburd

## Abstract

Describing the distribution of genetic variation across individuals is a fundamental goal of population genetics. In humans, traditional approaches for describing population genetic variation often rely on discrete genetic ancestry labels, which, despite their utility, can obscure the complex, multifaceted nature of human genetic history. These labels risk oversimplifying ancestry by ignoring its temporal depth and geographic continuity, and may therefore conflate notions of race, ethnicity, geography, and genetic ancestry. Here, we present a method that capitalizes on the rich genealogical information encoded in genomic tree sequences to infer the geographic locations of the shared ancestors of a sample of sequenced individuals. We use this method to infer the geographic history of genetic ancestry of a set of human genomes sampled from Europe, Asia, and Africa, accurately recovering major population movements on those continents. Our findings demonstrate the importance of defining the spatial-temporal context of genetic ancestry to describing human genetic variation and caution against the oversimplified interpretations of genetic data prevalent in contemporary discussions of race and ancestry.

## Introduction

The genomes of present-day individuals existed at every point in the past, scattered across geographic space and contained within the ancestors from whom they will eventually inherit their genetic material. The way those ancestors moved across space through history determines spatial patterns of genetic relatedness in the present (Bradburd and Ralph, 2019). Understanding these spatial patterns is vital both for the identification of the genomic basis of phenotypic variation (Battey et al., 2020) and for knowledge of the demographic history of a species (e.g. Ralph and Coop, 2013). Conversely, ignoring spatial demographic history can have serious implications for genome-wide association studies or the identification of loci involved in local adaptation (Zaidi and Mathieson, 2020).

Genetic variation in humans is often summarized with discrete geographic labels, but these can be inaccurate and misleading (Coop, 2022; Lewis et al., 2023). Even when loosely based on geographic history, genetic ancestry labels oversimplify a complex picture because they implicitly focus on only a single point in time.

For example, based on our current best understanding of human origins, all living individuals are “African” (regardless of the geography of their recent ancestors) on considering their ancestry ∼200ky before present. Advances in the study of ancient DNA have revealed a lack of genetic continuity within geographic regions (e.g. Haak et al., 2015; Allentoft et al., 2015; Racimo et al., 2020; Mattila et al., 2023), further highlighting the shortcomings of genetic ancestry labels. The fact that these labels are generated using statistical genetics approaches gives them the veneer of authenticity, further reifying problematic notions of race and ancestry in society (Lewis et al., 2023; Carlson et al., 2022).

At a technical level, many of the existing statistical methods for quantifying ancestry average over the ages of shared ancestors in the sample, effectively “flattening” the temporal component of the genealogy that connects all individuals within a species (Lawson et al., 2012; Pritchard et al., 2000). In reality, any pair of individuals is connected by many shared ancestors from whom each have inherited some portion of their genome (Mathieson and Scally, 2020). This flattening has the effect of painting a static notion of ancestry, rather than one that changes as it proceeds backwards in time.

If we knew the identities, locations, and ages of the ancestors of a sample, we could much more precisely and accurately report the geographic ancestry of a set of modern-day individuals through time. Moreover, we could learn about the history of dispersal in a species, identifying major population movements, demographic events, and barriers to migration. Although such detailed pedigree information is rarely available (but see Aguillon et al. (2017) and Anderson-Trocmé et al. (2023)), it nevertheless *is* possible to learn about the pedigree ancestors that are *shared* among individuals in a sample. This is because genetic relationships between samples, as well as the identities of the shared ancestors via whom they are related, are encoded in an interwoven collection of gene genealogies called an ancestral recombination graph (ARG) (Griffiths and Marjoram, 1997; Lewanski et al., 2024; Wong et al., 2023).

Recent advances in statistical and computational population genetics (Speidel et al., 2019; Kelleher et al., 2019; Wohns et al., 2022; Deng et al., 2024) have facilitated the inference of an ARG from large numbers of whole genomes. The ARG is a record of all coalescence and recombination events since the divergence of the sequences under study, and therefore specifies a complete genealogy of the sample at each genomic position. This record can be represented as a tree sequence (Kelleher et al., 2016, 2018) – an ordered set of trees, localized to adjacent regions of the genome, describing the gene genealogies of a set of samples at every genomic position. Each internal node in these local genealogies represents a haplotype within an ancestor from whom two or more sampled individuals have co-inherited a portion of their genome. By determining where and when each of these ancestors lived, we can, in principle, reconstruct the geographic history of a set of modern day individuals, documenting the path through space and time by which their genomes came to them. For example, Wohns et al. (2022) introduced a nonparametric approach that estimates ancestor locations by successively averaging the coordinates of sample locations in a postorder traversal of the ARG to their most recent common ancestor. In addition, Osmond and Coop (2021) and Deraje et al. (2024) describe likelihood methods for locating genetic ancestors and estimating migration rates based on a model of branching Brownian motion utilizing either samples of local gene trees or the full ARG.

Here, we present and validate an additional method for achieving this goal. Our method, called gaia (geographic ancestor inference algorithm), efficiently infers the geographic locations of the shared ancestors of a modern, georeferenced sample. We apply gaia to a tree sequence of humans sampled in Europe, Asia, the Middle East, and Africa (Wong et al., 2023), and are able to reconstruct the geographic history of human ancestry over the last two million years.

## Results

### Inferring the locations of shared ancestors

`Gaia` works by fitting a minimum migration cost function to each genomic position in an ancestral haplotype using the generalized parsimony algorithm and the local gene genealogy relating the sampled genomes at the given position (Fig. 1). Because the neighboring gene genealogies in a tree sequence are highly correlated, we are able to efficiently maintain the state of parsimony calculations as we iterate over the local genealogies in a tree sequence that contain the ancestral haplotype. Once these cost functions are computed for all genomic positions, we average them and assign the ancestral haplotype to the geographic location that minimizes its average cost function. These assignments then have a straightforward interpretation: they correspond to the geographic location that minimizes the overall migration cost of an average ancestral base pair. In our implementation, migration cost is a function of geographic distance, and the overall migration cost in a local genealogy is simply the sum of all ancestor-descendant migration costs.

**Figure 1.**
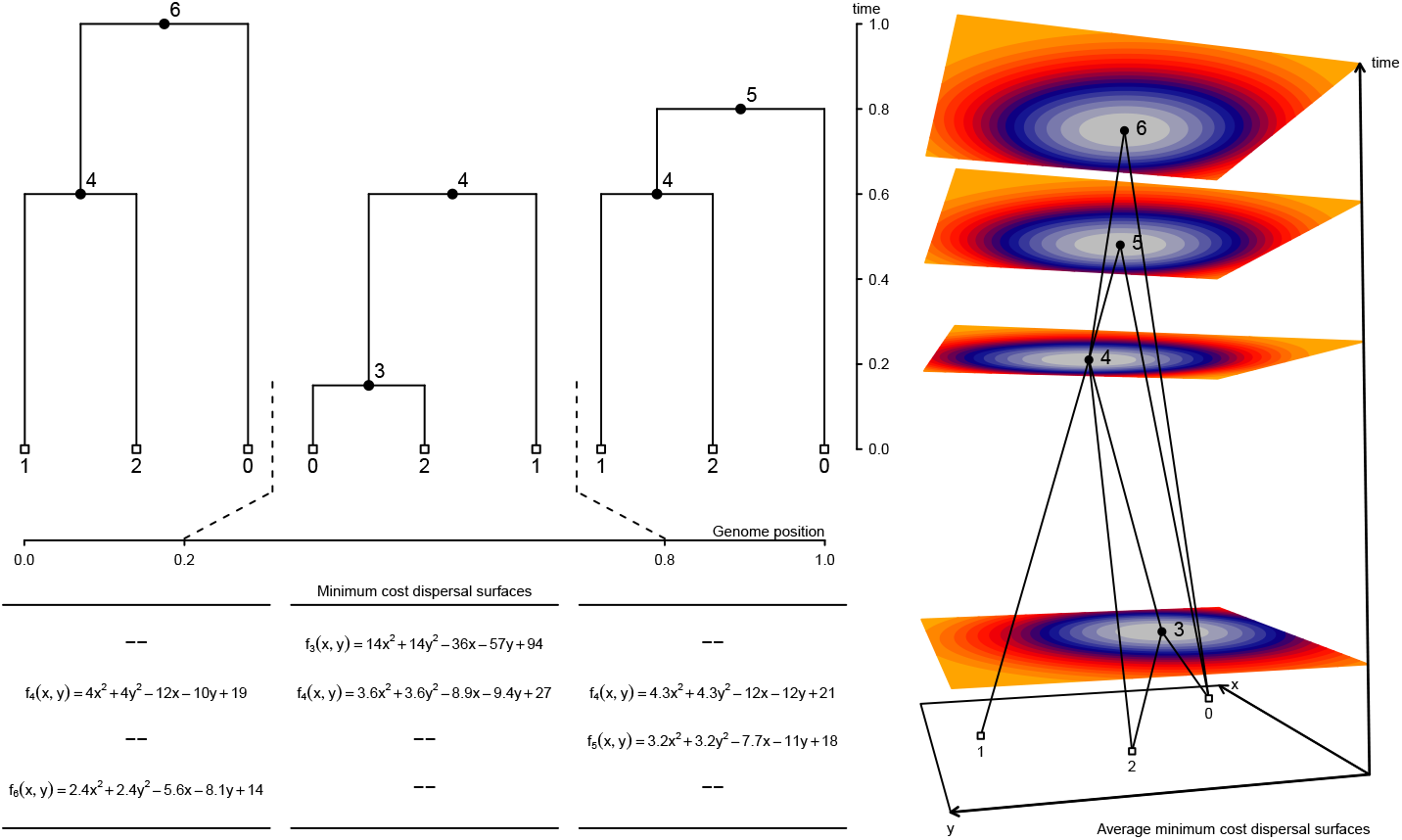
Conceptual overview of method. For each local tree we use the dynamic programming method of Sankoff and Rousseau (1975) to fit a minimum cost dispersal surface to the genealogical relationships of the georeferenced sample nodes. In this example, we use squared Euclidean distance as the cost function and *f*_*u*_(*x, y*) returns the smallest sum of squared dispersal distances between all ancestor-descendant node pairs that can be obtained when node *u* is at location (*x, y*). Using the genomic spans of local trees as weights, we then take a weighted average of local surfaces to assign each node a single average minimum cost dispersal surface. Here, node 4 appears in all three local trees and its final fitted surface is the weighted average of the three local surfaces. By contrast, nodes 3, 5, and 6 appear in a single local tree and their final fitted surfaces are identical to the surface in the local tree in which they appear. The perspective plot in the rightmost panel displays the ancestral recombination graph encoding the local trees along with the final fitted surface for each node. Non-sample nodes are positioned at the minimum cost point on the surface (warmer colors denote higher costs).

To validate the performance of gaia for inferring the geographic locations of nodes in a tree sequence, we simulated genetic data under different spatial models using SLiM (Haller and Messer, 2023). Gaia performs well under both expansionary and stationary demographic histories and over a range of dispersal kernels (both magnitude and shape) (Figs. S3,S4). We also demonstrate that we can use the reconstructed geographic distances between nodes in the tree sequence to estimate the parent-offspring dispersal distance for both Gaussian and long-tailed dispersal kernels (Figs. S1,S2).

### Tracking human ancestors through space and time

We inferred the geographic locations of ancestors of a contemporary sample of 2140 georeferenced human genomes from the Human Genome Diversity Project (Cann et al., 2002; Li et al., 2008) using a dated tree sequence inferred for chromosome 18 by Wohns et al. (2022) (Fig. 3). We focus on ancestry of the subset of individuals sampled from the continents of Europe, Asia, and Africa, consisting of 1070 contemporary individuals. The tree sequence for these individuals consists of 28,154 local genealogies, containing 114,606 ancestral nodes and spanning approximately 80,000 generations of human history. An equal area discrete global grid (Barnes and Sahr, 2023) (cell spacing approximately 800 km) intersected with Earth’s landmass provided a set of habitable areas, and individual sample locations were assigned to the nearest grid cell. Although a variety of complex migration cost matrices can be envisioned, including ones that incorporate long distance dispersal as well as temporal variation in the set of habitable areas, we opted for the simplest model that only allows migration between neighboring grid cells and assigns a unit cost to each migration event. With these assumptions, we used gaia to locate ancestral nodes to the grid cells with the lowest migration cost. For many ancestral nodes, especially older nodes, multiple grid cells may be optimal or near-optimal (Fig. 2). Because our summaries ignore near-optimal solutions, they do not explore the full range of uncertainty in ancestral locations and should be viewed not as precise statements on where ancestors lived and how they moved, but rather as summaries of major trends.

**Figure 2.**
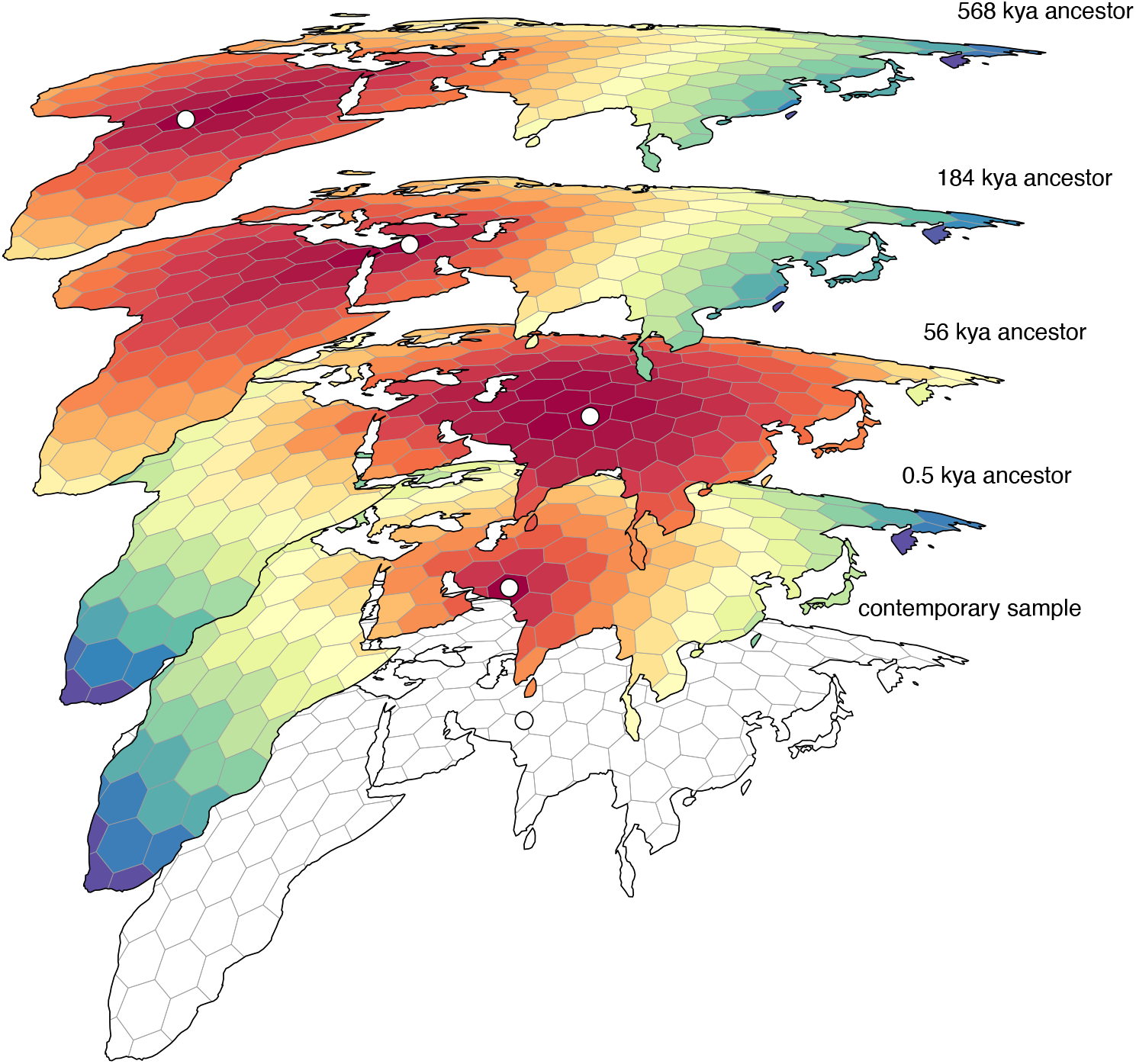
An example set of geographically referenced genetic ancestors of a contemporary sample from Asia. Colored grid cells denote the migration costs calculated by gaia (warmer colors denote lower costs) and each ancestor (denoted by a white point) is positioned within the minimum cost grid cell. For most ancestors there are many near-optimal locations that are ignored by our summary methods, which should therefore be viewed as summarizing major trends rather than as precise statements on where ancestors lived and how they moved.

**Figure 3.**
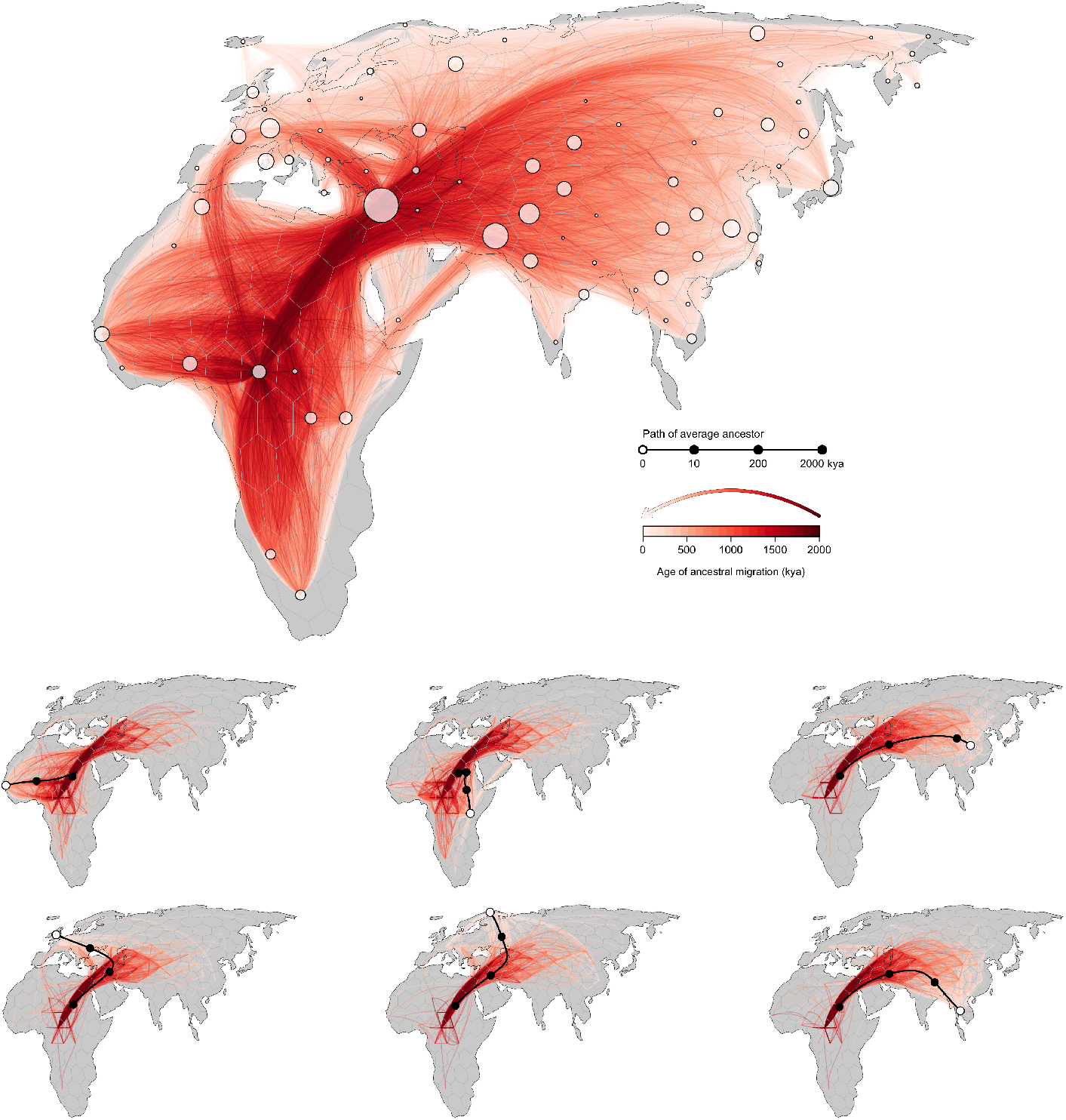
A georeferenced ancestral recombination graph (ARG). Red lines trace the inferred historical migrations of genetic ancestors of the sample (white points), with darker shading used to indicate movements that took place in the more distant past. Six contemporary samples are highlighted in the lower panels; black lines trace the average position of their genetic ancestors back through time, while red lines denote the subset of edges in the ARG that are ancestral to the samples.

Our inferred geographic chronology of the ancestors of the sample largely reconstructs major population movements in human prehistory, including the out-of-Africa expansion and the peopling of Eurasia (Fig. 4). Our estimates place some genetic ancestors of the sample in Asia and the Middle East well before the earliest fossil evidence of human dispersal out of Africa. The oldest nodes in the tree sequence are estimated to have occurred roughly 2 million years before present, and a small minority of these are inferred to be located in Asia and the Middle East. A similar pattern was observed by Wohns et al. (2022) in their analysis of chromosome 20. Nonetheless, from ca. 2 million to 200ky before present, the average positions of genetic ancestors to the geographic subsets of samples from Europe, Asia, the Middle East, and Africa are all inferred to be in Africa. This coincides with the time period when the majority of genome positions in these geographic subsets trace their descent to an ancestor in Africa. Between 200kya and the present, the average location of ancestors to the geographic subsets of samples from Europe, Asia, and the Middle East diverge from one another and begin to move toward the average of position of samples from those regions, while the average position of ancestors to the African subset of samples remains in Africa (Fig. 5).

**Figure 4.**
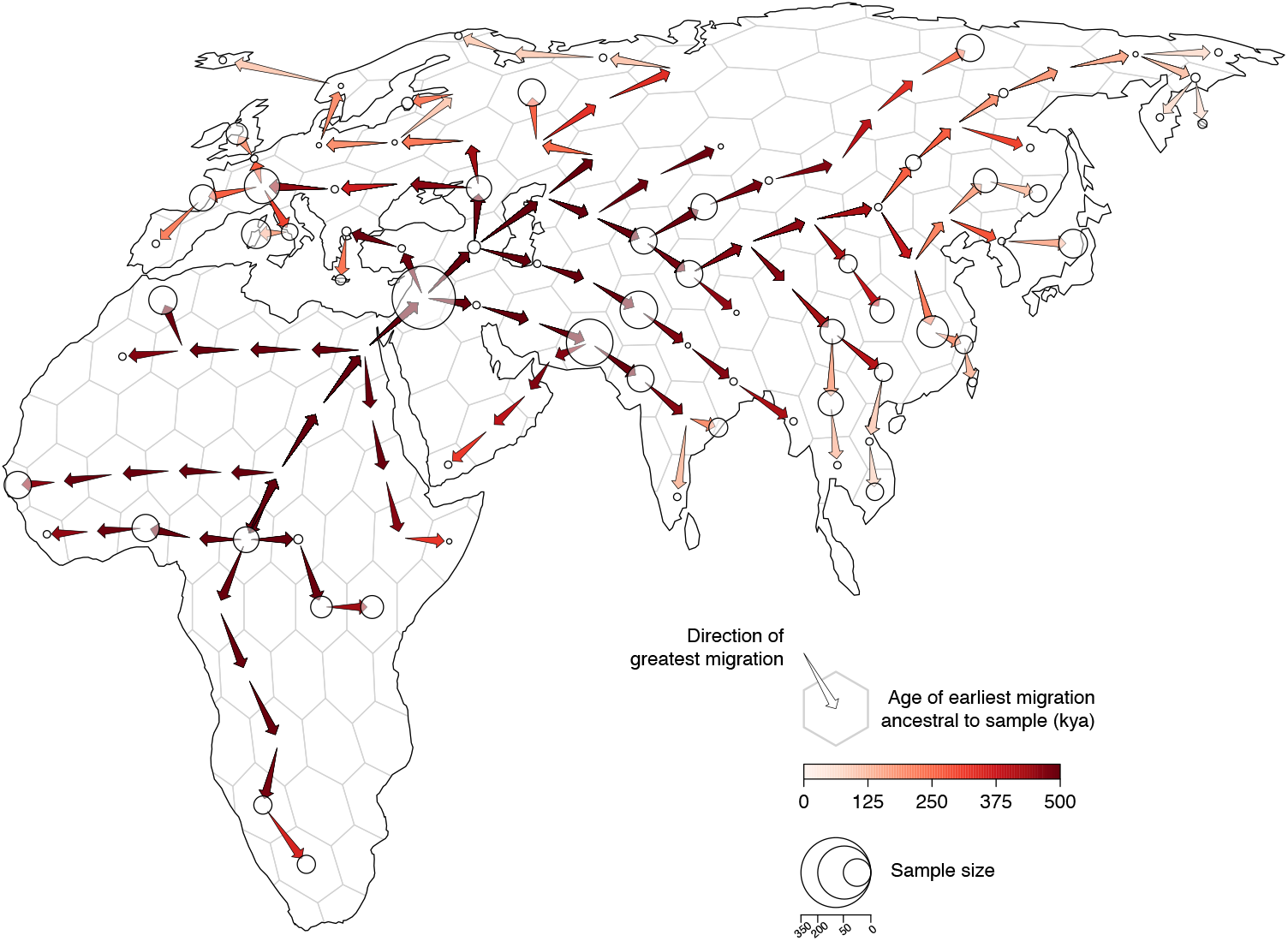
Geographic chronology of human genetic ancestry. Arrows point in the direction of greatest migration of the shared genetic ancestors of the sample and are colored according to the age of the earliest migration. Points show the distribution of sampled modern-day genomes. Point size is proportional to the number of samples from each locality.

**Figure 5.**
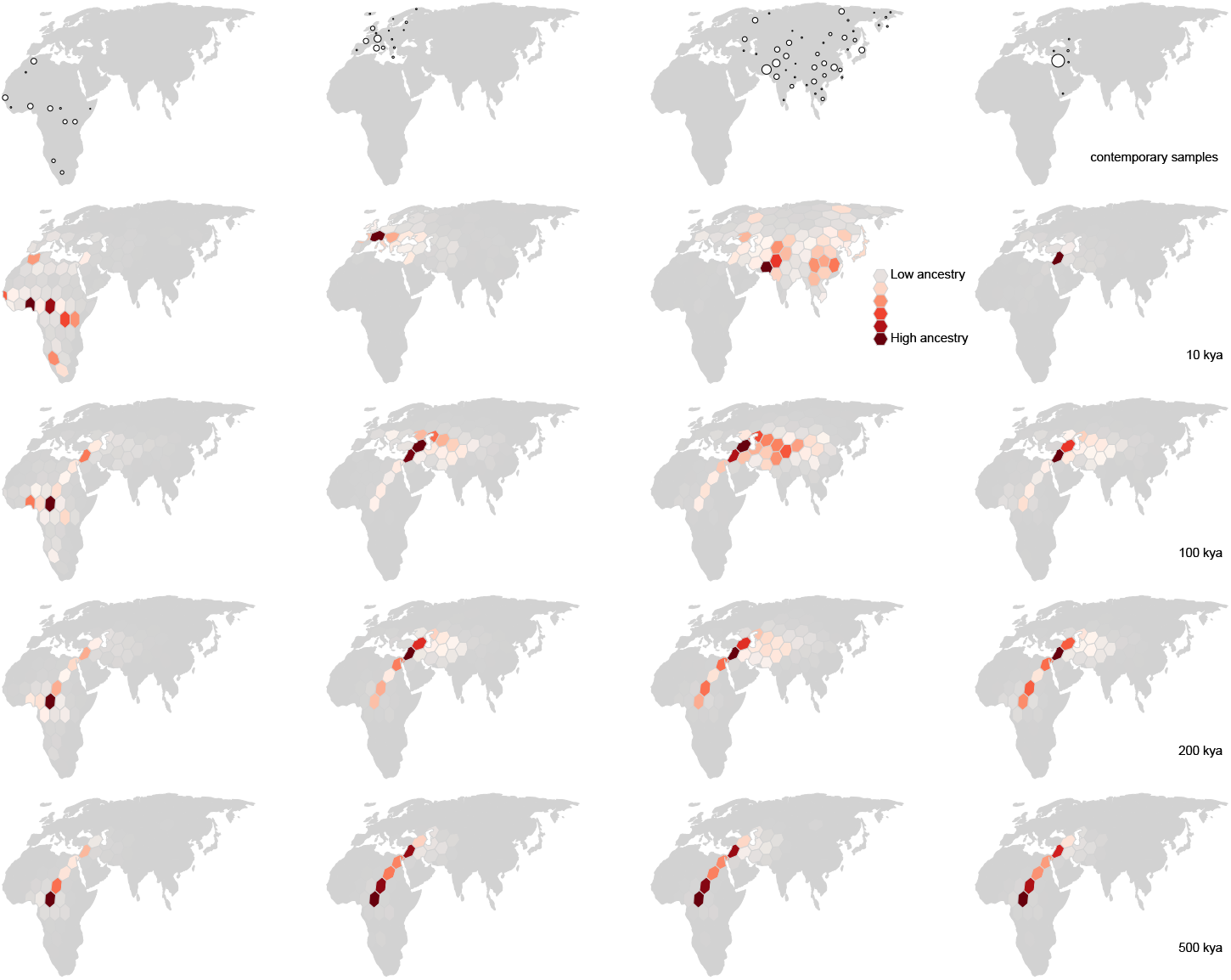
Inferred spatiotemporal ancestry coefficients through time. The fraction of genomic positions in each geographic subset of samples (top row) that trace ancestry to different geographic regions at different times in the past is represented by shades of red, with darker shading indicating greater ancestry (=a larger fraction). Point size is proportional to the number of sampled genomes from each locality.

### Geographic history of human ancestry

We use the georeferenced tree sequence to define a spatiotemporally explicit ancestry coefficient, which we then track across space and time to understand and quantify the genetic and geographic history of our sample. Only a subset of any given contemporary individual’s genome, *A*_*i*_(*t*) in individual *i*, is found in the ancestral nodes in the tree sequence at a given point in time *t* (once the local genealogy in a particular genomic region has coalesced, that portion of the genome is no longer represented in deeper sections of the ARG; therefore, the farther back in time we look, the less of any modern-day individual’s genome can be found). We define this spatiotemporally explicit ancestry coefficient, which we call *z*_*ik*_(*t*), as *A*_*ik*_(*t*)*/A*_*i*_(*t*): the proportion of individual *i*’s genome that exists in its ancestors in the ARG at time *t* and that is inherited from ancestors living within some prescribed geographic region *k*. This ancestry coefficient can be broadly understood as the proportion of a sample’s genome inherited from individuals living within a specified geographic region at a specific time.

Unlike ancestry labels, *z*_*ik*_(*t*) is *explicitly* associated with both a point in time and a region of space; we can therefore use it to understand how a sample’s ancestry has changed across space and time. At the present moment, the distribution of *z*_*ik*_ simply reflects the geography of modern-day sampling. By 100,000ya, we find that only 2.5% of modern-day samples are inheriting their genomes from ancestors in Europe (Fig. 5); instead, almost all ancestors contributing genomic material to the modern-day sample are inferred to have lived in Africa, Asia, and the Middle East. The spatiotemporal ancestry coefficients show similar trajectories for modern-day samples from Asia. During this same time interval, the proportion of sample ancestors found in Africa increases almost monotonically backward in time, from 16% at the present (again, reflecting the geography of modern-day sampling) to ∼90% by 1Mya. Interestingly, the proportion of the genomes of the sample inherited from ancestors inferred to be in Africa plateaus at ∼90%, with ∼10% of genome continuing to be inherited from ancestors in the Middle East along the eastern Mediterranean. (Note that this region contributed the largest number of samples to the dataset.)

### Large scale movement in human ancestry

To study large-scale movements of human ancestry, we introduce a new statistical summary of the georeferenced tree sequence: ancestry flux. Formally, if *A*_*i*_(*t*_*l*_,*t*_*r*_) is the amount of individual *i*’s genome that inherits from ancestors alive between [*t*_*l*_,*t*_*r*_) and *A*_*ijk*_(*t*_*l*_,*t*_*r*_) is the same amount that inherits from ancestors who moved from *j* to *k*, we define ancestry flux as 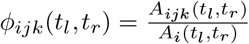: the proportion of *i*’s genome that exists in its ancestors in the ARG during the time period [*tl,t_r_*) and that is inherited from ancestors who moved from *j* to *k* during that same time period. This coefficient can be broadly understood as the proportion of a sample’s genome inherited from ancestors who moved between specified geographic regions during a particular time period in the past.

We discretized Europe, Asia, and Africa into equal-area hexagons (each approximately 800 sq km) and quantified ancestry flux between them in 2,500 year intervals between the present and 0.5Mya. We find consistent ancestry flux out of Africa into the Middle East during this time period, with a peak occurring between 100,000ya and 150,000ya (Fig. 6). Interestingly, nearly all ancestry flux out of Africa is estimated to occur through a northern route across the Sinai Peninsula rather than a southern route across the strait at Babel-Mandeb. Approximately 30% of genomic positions in the modern-day sample trace ancestry to a northerly migration out of Africa but only 0.1% trace ancestry to a southerly migration route out of Africa.

**Figure 6.**
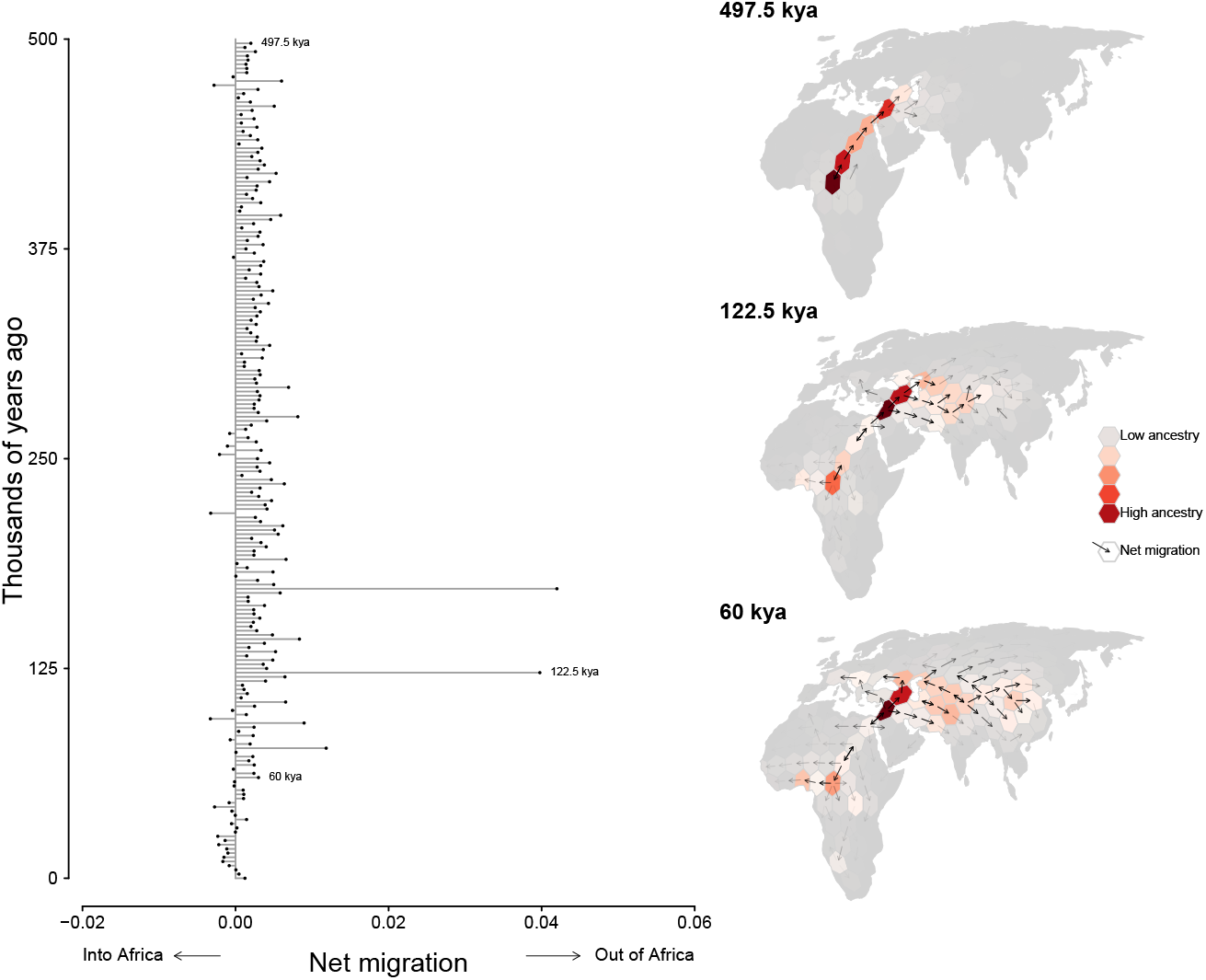
Inferred flux in spatiotemporal ancestry coefficients through time. Each bar depicts the net direction of historical migration of shared genetic ancestors of the modern-day sample. Inset maps highlight three different time intervals. Cells are colored by the fraction of genomic positions that find ancestry in them at that time and arrows depict the direction of greatest migration with opacity scaled by the magnitude migration.

## Discussion

We have described a simple heuristic approach for reconstructing the geographic genetic history of a sample, which can be used to explore the timing and geographic location of demographic events. Patterns of genetic diversity and relatedness between individuals are created by a complex interplay between evolutionary forces acting over the history a population. By uncovering these patterns, our method enables researchers to go beyond the standard practice of grouping individuals into a small number of static and ill-defined “populations.” Although this practice may be useful in some limited scenarios (e.g., for conservation and management), our results illustrate that it discards an immense amount of additional information that can be extracted from the data. Furthermore, when applied to humans, population assignment has the potential to lead to greater harm and confusion by conflating race, ethnicity, geography, and ancestry (Coop, 2022). A spatiotemporal summary of ancestors offers an intuitive and informative path toward understanding ancestry, demography, and major population movements through time.

Of course, the geographic histories inferred by gaia are not perfectly accurate; individual dispersal decisions are the result of countless external factors that we cannot hope to capture using a simple heuristic model. For example, gaia assumes that lineages disperse independently of one another and that the dispersal process is independent of the coalescent process in a reconstructed gene tree; both assumptions are likely to be violated in empirical datasets. Moreover, errors in the tree sequences inferred by upstream tools such as relate (Speidel et al., 2019) and tsinfer (Kelleher et al., 2019) have the potential to propagate into the output of gaia. Additionally, the sampling process itself may impact our results, insofar as the accuracy of geographic inference of ancestor locations depends on the distribution of sampled individuals relative to that of their ancestors. Finally, we caution that gaia sheds light on the geographic and temporal distribution of *shared genetic ancestors*; even if we had perfect information on the geographic locations of all ancestors, it would not necessarily inform our understanding of the geographic distribution of the general census population alive at any point in the past, though the two are likely correlated. As with all population genetic approaches, historical individuals that contributed no ancestry to the modern-day sample remain inscrutable.

Despite these limitations, it is clear that the ability to infer the locations of ancestors in the tree sequence opens exciting avenues of future research. Simply summarizing the georeferenced tree sequence has the potential to yield valuable insights into the evolutionary and natural history of a population, including identifying barriers to dispersal, shifts in dispersal regimes (magnitude and direction) through time, and the geography of ecological dynamics (at least with respect to genetic ancestry). A straightforward direction for extending gaia would be to explicitly incorporate some of these elements – e.g., heterogeneous dispersal rates between different regions. As the transitions between different geographic states occur on an arbitrary network (generated, in the applications presented here, via geographic proximity), we can easily incorporate different geographies; for example, dispersal routes that connect far-flung regions, which may be of interest when species occasionally experience human-mediated dispersal (e.g., (Zhan et al., 2023)). More broadly, the ability to study the geography of genealogies heralds an exciting growth in the ability of the field of population genetics to shed light on population ecological processes governing the movement, distribution, and density of individuals across space and through time.

## Methods

### Main approach: computing minimum migration fits to local trees

Given a set of sampled genomes with known locations and a tree sequence that relates the samples, gaia efficiently reconstructs the locations of genetic ancestors using a minimum migration heuristic. Conceptually, gaia may be thought of as proceeding in three steps (Fig. 1).

In the first step, a minimum migration cost function *f*_*uk*_ is fitted to each node *u* in each local tree *k* in which *u* appears using the generalized parsimony algorithm (Sankoff, 1975; Sankoff and Rousseau, 1975). For each spatial location, *f*_*uk*_(*x*) then returns the smallest sum of migration costs between all ancestor-descendant pairs in local tree *k* that can explain the sampled locations when *u* is at location *x*. We assume migration cost is a function of migration distance and provide three cost function implementations. In each of these three cases we allow migration distances to optionally be weighted by inverse branch length. For a continuous state space, we use either the squared Euclidean distance or the Manhattan distance. Squared Euclidean distance leads to cost functions that are convex quadratic functions of the spatial locations (Maddison, 1991); Manhattan distance, to convex piecewise linear functions (Csűrös, 2008). For a finite set of geographic locations, a square matrix with arbitrary transition costs between all pairs of geographic states leads to minimum migration costs that are discrete functions defined on the set of geographic locations (Clemente et al., 2009).

In the second step, we form a new function *F*_*u*_ by taking a weighted average of *f*_*uk*_ over the local trees in which *u* appears, using the genomic spans of each local tree as weights. We interpret *F*_*u*_ as the minimum migration cost of an average ancestral base pair. In other words, if one were to take each sampled base pair that inherits from *u* and fit a minimum migration cost function to its coalescent history, the average of those would look like *F*_*u*_. In the Appendix, we present a detailed description of the algorithm we use for computing *F*_*u*_ from the local tree fits *f*_*uk*_. Considerable computational savings can be achieved by using the succinct tree sequence encoding (Kelleher et al., 2016), which allows us to efficiently maintain the state of parsimony calculations as we iterate over local trees. As detailed in the Appendix, the local tree fits also allow us to compute an estimate of effective migration rate, which we define as the mean lineage migration cost of an average base pair. That is, if one were to take the coalescent history of each sampled base pair and compute the mean the per-branch migration cost in a most parsimonious migration history, the average of those – taken over all sampled base pairs – would equal the effective migration rate.

Finally, we locate each ancestor by finding the spatial location *x* where *F*_*u*_(*x*) is smallest. A constrained (rather than global) minimization of *F*_*u*_ could also be used to locate ancestors if some spatial locations are known to be uninhabitable. Although we do not pursue this possibility in the current study, uncertainty in ancestor locations could be explored by sampling from exp(–*λ*[*F*_*u*_(*x*) –min_*z*_ *F*_*u*_(*z*)]), where *λ* is a tuning parameter that controls how strongly deviations from the spatial location that minimizes the overall migration cost are penalized.

### Spatiotemporal ancestry coefficients

We use our ancestor location estimates to define two spatially and temporally explicit ancestry coefficients as follows. In the first case, consider a subset *i* of the sampled genomes. Define *A*_*i*_(*t*) to be the total amount of sample material in the subset that is descended from ancestral material at a point in time *t* in the past. Define *A*_*ik*_(*t*) to be the total amount of sample material descended from ancestral material in geographic location *k* at time *t* in the past. We then define the spatiotemporal ancestry coefficient for subset *i* as 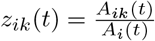, the fraction of sample material that is descended from ancestral material in geographic location *k* at time *t* in the past. Thus, if we were to pool all sample material in subset *i* whose coalescent history extended at least *t* time units into the past, *z*_*ik*_(*t*) is the probability that we can trace the coalescent history of a randomly selected base pair from that pool to an ancestral base pair in geographic location *k* at time *t* in the past.

Next, define *A*_*i*_(*t*_*l*_,*t*_*r*_) to be the total amount of sample material in the subset that is descended from ancestral material present during an interval in time [*t*_*l*_,*t*_*r*_) in the past. Define *A*_*ijk*_(*t*_*l*_,*t*_*r*_) to be the total amount of sample material descended from ancestral material that migrated from geographic location *j* to location *k* during an interval in time [*t*_*l*_,*t*_*r*_) in the past (where *j* is not equal to *k*). We then define the spatiotemporal ancestry flux coefficient for subset *i* as 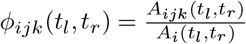. Thus, if we were to pool all sample material in subset *i* whose coalescent history extended at least [*t*_*l*_,*t*_*r*_) time units into the past, *ϕ*_*ijk*_(*t*_*l*_,*t*_*r*_) is the probabily that we can trace the coalescent history of a randomly selected base pair from that pool to an ancestral base pair that migrated from geographic location *j* to location *k* during the interval [*t*_*l*_,*t*_*r*_) in the past.

As defined, our spatiotemporal ancestry coefficients require knowing the geographic locations of ancestral lineages at arbitrary points in time in the past but our minimum migration location estimates (i.e., argmin_*x*_ *F*_*u*_(*x*)) are available only for the two endpoints of an ancestral lineage. By using the migration cost function and conditioning on these endpoint states, however, we can sample a minimum cost migration history for each lineage and interpolate the history between the endpoint times of the ancestral lineage. In this way we can locate ancestors at arbitrary points in time. Further details about this procedure are presented in the Appendix.

### Simulation study: testing ancestor location accuracy

We tested performance of gaia using spatially explicit forward-time simulations in SLiM v4.0.1 (Haller and Messer, 2023). Simulated individuals were semelparous and hermaphroditic and diploid for a single chromosome with 10^8^ basepairs and a recombination rate of 10^−8^ per basepair per generation. Individuals coexisted on a two-dimensional square plane with reflecting boundaries and a side length of 1000*σ* units, where *σ* was the standard deviation of the dispersal kernel.

Population regulation occurred via density dependent effects on fecundity and survival. The number of offspring produced by a focal individual in generation *t* was Poisson-distributed with a mean equal to 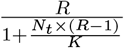, where *N*_*t*_ is the local population density in a circle of radius 3*σ* around the focal individual at generation *t, R* is the population growth rate at low density, and *K* is the local carrying capacity density. During reproduction, each individual (the “mother”) chose a mate (the “father”) uniformly at random from the set of individuals living within a radius of 3*σ* from itself. A random location centered on the mother’s location was then chosen for each offspring by drawing from a dispersal kernel with a standard deviation of *σ*. Each offspring produced in generation *t* survived to reproduce in generation *t* + 1 with probability 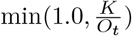, where *O*_*t*_ is the local population density in a circle of radius 3*σ* around the focal offspring in generation *t*.

We conducted simulations using either a Gaussian dispersal kernel or a double exponential dispersal kernel and 10 different *σ* levels that were equally spaced from 0.2 to 2.0. The Gaussian kernel emulates a Brownian motion while the double exponential kernel emulates a Laplace motion, which generates more extreme displacements of offspring relative to Brownian motion.

We conducted 10 replicate simulations for each *σ* level under each dispersal kernel. For all simulations, we set *R* = 2 and 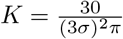. Each simulation was initiated from a randomly located founder population of 30 individuals that was allowed to grow and disperse for 10,000 generations, after which the simulation was terminated and the recorded (simplified) tree sequence of extant individuals was saved for analysis.

Simulations conducted under the Gaussian dispersal kernel were analyzed using squared Euclidean distance weighted by inverse branch length as the transition cost function. Simulations conducted under the double exponential dispersal kernel were analyzed using Manhattan distance weighted by inverse branch length as the transition cost function. For all simulations we evaluated our ability to accurately estimate effective migration rates and the locations of genetic ancestors. We defined the true effective migration rate (*σ*_*e*_) to be the maximum likelihood estimate of *σ* under each dispersal kernel assuming we had complete knowledge of the locations of all ancestral individuals in the tree sequence.

### Empirical example: reconstructing human migration out of Africa

We applied gaia to a contemporary sample of georeferenced human genomes from the Human Genome Diversity Project using the dated tree sequence inferred for chromosome 18 by Wohns et al. (2022). Prior to analysis, we first simplified the tree sequence to include only individuals from Africa, Asia, and Europe. Certain non-sample nodes in the tree sequence are clear outliers with respect to the number of edges in which they participate. For example, one node has nearly 3,000 immediate descendants, well above the median number of 6. As these genealogical outliers have the potential to strongly bias geographic reconstructions, we further simplified the tree sequence by removing edges to or from non-sample nodes in the upper two percent of indegree or out-degree counts. The resulting simplified tree sequence consisted of 28,154 local genealogies, containing 114,606 ancestral nodes and spanning approximately 80,000 generations of human history.

To explore historical geographic patterns in human genetic ancestry over the last one-half million years we created a set of equal-interval time bins spanning 100 generations (or 2,500 years using a 25 year generation time) that extended from the present to 20,000 generations in the past. An equal area discrete global grid (Barnes and Sahr, 2023) (cell spacing approximately 800 km) intersected with Earth’s landmass provided a set of habitable locations. Individual samples were then mapped to the nearest grid cell and we used gaia to compute *F*_*u*_ for each node in the empirical tree sequence using a simple cost matrix that assigned a unit cost to each migration event and only allowed migration between neighboring grid cells.

Each internal node in the tree sequence was then geographically referenced by sampling uniformly at random one of the grid cells that minimized its migration cost function *F*_*u*_ (for some nodes there may be a unique grid cell that achieves this minimum but for others there may be multiple). This procedure was repeated 100 times and spatiotemporal ancestry coefficients were computed for each realization and then averaged.

## Acknowledgements

We thank Yaniv Brandvain, Jed Carlson, Graham Coop, Puneeth Deraje, Doc Edge, James Kitchens, Marjorie Weber, and members of the Bradburd lab for comments and discussion that helped improve the research presented here. This research was supported by National Science Foundation (award number DMS-2052653), the National Institute of General Medical Sciences of the National Institutes of Health under award numbers R35GM151145 (JT) and R35GM137919 (GB). The content is solely the responsibility of the authors and does not necessarily represent the official views of the National Institutes of Health.

## Supplementary Information

## Appendix

### A1. Overview

We first review generalized parsimony as it applies to a single gene tree (Sankoff, 1975; Sankoff and Rousseau, 1975). For each node *u* we maintain three cost functions. The *node cost* function *g*_*u*_(*x*) assigns a cost to each geographic state *x* (when *u* is in state *x*) that gives the minimum migration cost required to explain the geographic states of all sample nodes whose most recent common ancestor is *u*. If *u* is a sample node, we set *g*_*u*_(*x*) = 0 if *x* = *x*_*u*_ and *g*_*u*_(*x*) = ∞ if *x* ≠ *x*_*u*_. When *u* is not a sample node, *g*_*u*_ is formed by the sum of the *stem cost* functions of its children.

The stem cost function for a node *u* is given by *h*_*u*_(*x*) = min_*z*_ [Δ_*u*_(*x, z*) + *g*_*u*_(*z*)]. The function Δ_*u*_(*x, z*) assigns a cost to the migration from state *x* to state *z* over the branch leading to node *u*. Here, we assume that Δ_*u*_ can be factored as: Δ_*u*_(*x, z*) = *ϕ*(*τ*_*u*_) * Δ(*x, z*), where *τ*_*u*_ is the length of the branch leading to node *u* and Δ is a time-independent transition cost function. Typical choices for the time-dependent component are 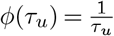 or *ϕ*(*τ*_*u*_) = 1, the latter making the whole cost function independent of branch lengths.

The node cost and stem cost functions can be computed in a single post-order tree traversal. A second pre-order tree traversal can then be used to compute the *final cost* functions for each node. The final cost *f*_*u*_(*x*) gives the minimum migration cost required to explain the geographic states of *all* sample nodes when *u* is in state *x*. When *u* is the root of the tree, *f*_*u*_(*x*) = *g*_*u*_(*x*). Otherwise, 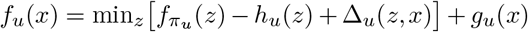, where *π*_*u*_ represents the immediate ancestor of node *u*.

### A2. Algorithm

Here we describe how we use the succinct tree sequence encoding (Kelleher et al., 2016) to efficiently compute node cost and stem cost functions for each local genealogy. Under the coalescent with recombination, nearby gene trees are highly correlated. Moving between adjacent local trees typically requires only a small number of subtree-prune-and-regraft (SPR) operations. By keeping an index of the edges involved in these SPR operations and the order in which they need to be applied, the succinct tree sequence encoding allows us to efficiently maintain the state of parsimony calculations as we iterate over the local genealogies.

A detailed description of the tree sequence data structure can be found in Kelleher et al. (2016). For our purposes, it suffices to know that relationships among all *N* nodes in the tree sequence are recorded in the edge table *E*. Each row in the edge table records an ancestor-descendant relationship, and the indices of nodes involved in that relationship can be accessed as *E*[*k*].parent and *E*[*k*].child. The relationship encoded by an edge applies to the half-open genomic interval between *E*[*k*].left (inclusive) and *E*[*k*].right (exclusive). As we move along the genome from left to right, the index vectors *I* and *O* give the insertion and removal order of the edges needed to build each local gene tree topology (recorded in the vector *π*). The following algorithm is based on the branch statistic algorithm that appears in Ralph et al. (2020) but differs in its treatment of sample weights and how they are propagated along a genealogy.

#### Algorithm P (generalized parsimony)

Given a set of georeferenced samples related by a tree sequence with length *L*, compute the genome-wide average final cost function *F*_*u*_(*x*) and the genome-wide average migration rate statistic *σ*_*P*_. The function *F*_*u*_(*x*) returns the average minimum migration cost required to explain the observed geographic states of all sample nodes when *u* is in state *x*, where the average is taken over all local genealogies (weighted by their genomic span) where node *u* appears. The statistic *σ*_*P*_ is the average per-branch migration cost in a most parsimonious migration history averaged over all local genealogies weighted by their genomic span.

**S1**. [Initialization.] For 0 ≤ *u* < *N* set *π*_*u*_ ← −1, *s*_*u*_ ← 0, *g*_*u*_(*x*) ← 0, *F*_*u*_(*x*) ← 0. Then, if *u* is a sample node and *x* ≠ *x*_*u*_, set *g*_*u*_(*x*) ← ∞ and *F*_*u*_(*x*) ← ∞

Finally, set *j* ← 0, *k* ← 0, *σ*_*P*_ ← 0, *t*_*l*_ ← 0, *s*_*P*_ ← 0.

**S2**. [Terminate.] If *j* = |*E*| terminate.

**S3**. [Edge removal loop.] If *k* = |*E*| or *t*_*l*_ ≠ *E*[*O*_*k*_].`right` go to S6.

**S4**. [Remove edge.] Set *u* ← *E*[*O*_*k*_].`parent`, *v* ← *E*[*O*_*k*_].`child`, *w* ← *π*_*u*_, and *k* ← *k* + 1. Then, if *w* ≠ −1, set *g*_*w*_(*x*) ← *g*_*w*_(*x*) −*h*_*u*_(*x*). Finally, set *g*_*u*_(*x*) ← *g*_*u*_(*x*) −*h*_*v*_(*x*), *π*_*v*_ = −1, *v* ← *u, u* ← *w*.

**S5**. [Update node and stem costs.] While *u* ≠ −1, set *w* ← *π*_*u*_ and if *w* ≠ −1 set *g*_*w*_(*x*) ← *g*_*w*_(*x*) −*h*_*u*_(*x*). Then set *h*_*v*_(*x*) ← min_*z*_ [Δ_*v*_(*x, z*) + *g*_*v*_(*z*)], *g*_*u*_(*x*) ← *g*_*u*_(*x*) + *h*_*v*_(*x*), *v* ← *u, u* ← *w*. Afterward, go to S3.

**S6**. [Edge insertion loop.] If *j* = |*E*| or *t*_*l*_ ≠ *E*[*I*_*j*_].`left` go to S9.

**S7**. [Insert edge.] Set *u* ← *E*[*I*_*j*_].`parent`, *v* ← *E*[*I*_*j*_].child, *w*← *π*_*u*_, and *j* ← *j* + 1. Then, if *w* ≠ −1, set *g*_*w*_(*x*) ← *g*_*w*_(*x*) − *h*_*u*_(*x*). Then set *h*_*v*_(*x*) ← min_*z*_ [Δ_*v*_(*x, z*) + *g*_*v*_(*z*)]. Finally, set *g*_*u*_(*x*) ← *g*_*u*_(*x*) + *h*_*v*_(*x*), *π*_*v*_ = *u, v* ← *u, u* ← *w*.

**S8**. [Update node a nd stem costs.] While *u* ≠ −1, set *w* ← *π*_*u*_ and if *w* ≠ −1 set *g*_*w*_(*x*) ← *g*_*w*_(*x*) −*h*_*u*_(*x*). Then set *h*_*v*_(*x*) ← min_*z*_ [Δ_*v*_(*x, z*) + *g*_*v*_(*z*)], *g*_*u*_(*x*) ← *g*_*u*_(*x*) + *h*_*v*_(*x*), *v* ← *u, u* ← *w*. Afterward, go to S6.

**S9**. [Genomic span of tree.] Set *t*_*r*_ ← *L*. If *j* < |*E*| set *t*_*r*_ ← min(*t*_*r*_, *E*[*I*_*j*_].left). Then, if *k* < |*E*| set *t*_*r*_ ←min (*t*_*r*_, *E*[*O*_*k*_].`right`. Set *s* ← *t*_*r*_ −*t*_*l*_.

**S10**. [Update average migration costs.] Set *σ* ← 0, *n* ← 0, *s*_*P*_ ← *s* + *s*_*P*_. Then visit each node *u* in a pre-order traversal and set *s*_*u*_ ← *s*_*u*_ +*s*. If *π*_*u*_ = −1 set *f*_*u*_(*x*) ← *g*_*u*_(*x*) and set *σ* ← *σ* +min_*z*_ *f*_*u*_(*z*); otherwise, set 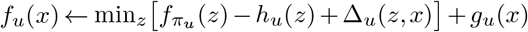 and set *n* ← *n* + 1. Then, set 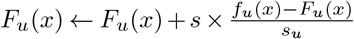.

**S11**. [Update average migration rate.] Set 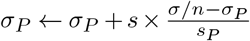.

**S12**. [Tree loop tail.] Set *t*_*l*_ ← *t*_*r*_. Go to S2.

We begin in S1 by initializing the cost functions for each node to zero except in the case of sample nodes, for which we set the cost function to positive infinity for all states not equal to the observed state. We then set the parent of each node to −1 (signifying the null element) so that the initial state of the tree sequence is a forest of disconnected nodes. The average cost functions (denoted by the corresponding capital letters) are also initialized to 0 as these will be updated as we iterate over the tree sequence.

The meat of the algorithm occurs in steps S4 and S5 of the edge removal loop and in steps S7 and S8 of the edge insertion loop. Removal of an edge from node *u* to *v* will alter the node and stem cost functions along the path from *u* back to the root of the genealogy. Step S4 prepares for this by first subtracting the stem costs of *u* and *v* from the node costs of their respective parents. In step S5, we walk back along the path from *u*’s parent to the root and recompute new stem costs and node costs given the updated node costs at the head of the path. Insertion of an edge from node *u* to *v* will similarly alter the node and stem cost functions along the path from *u* back to the root of the genealogy. Step S7 prepares for this by first substracting the stem cost of *u* from the node cost of its parent and then computing the stem cost of *v* and adding it to the node cost of *u*. In step S8, we walk back along the path from *u*’s parent to the root as before and recompute new stem costs and node costs given the updated node costs at the head of the path.

Upon reaching step S9, we have finished constructing the tree together with its node cost and stem cost functions. We record the genomic span of the tree in the variable *s*, which will be the weight applied to the current tree in the weighted average. In S10, we perform a preorder traversal of the newly constructed tree and compute the final cost function for each node. At the same time, we increment the total weight *s*_*u*_ of each node by *s* and update the weighted averages *F*_*u*_(*x*) with the cost functions for the current tree. When we begin the traversal at the root(s) of the tree we also record the minimum migration cost *σ* needed to explain the sample distribution. We use this cost in S11, together with the number *n* of edges in the tree, to update the weighted average migration rate statistic *σ*_*P*_.

Correctness of the algorithm requires that edges are removed in order of nondecreasing right genomic coordinate and decreasing time and that edges are inserted in order of nondecreasing left genomic coordinate and increasing time (time is measured backward from the present). We assume that the index vectors *I* and *O* are constructed to satisfy these conditions; these assumptions form the basis of the conditional checks in steps S3 and S6 for determining when to exit the edge removal and insertion loops.

#### Relation to existing work

Several existing approaches to geographic inference with tree sequences merit discussion in relation to our own work. Wohns et al. (2022) introduced a nonparametric approach that estimates ancestor locations by successively averaging the coordinates of sample locations in a postorder traversal of the ARG to their most recent common ancestor. The resulting estimates are local estimates in the sense that the inferred location of an ancestor depends only on the locations of samples that trace some portion of their ancestry to that ancestor and on the topology of the corresponding subset of the ARG. By contrast, our approach estimates the location of an ancestor using information from all samples. Because all samples share common ancestry at some time in the past, even those samples that are not direct descendants of an ancestor can be informative about that ancestor’s location.

Osmond and Coop (2021) describe a likelihood method for locating genetic ancestors and estimating migration rates that is based on a model of branching Brownian motion. Their approach also uses information from all samples to estimate ancestral locations and can optionally estimate separate migration rates for deep and shallow time horizons. Unlike our method, inference is carried out on a sample of widely spaced genealogies rather than on the full tree sequence. We note that when squared Euclidean distance weighted by inverse branch length is used as the transition cost function, the maximum parsimony reconstruction on each marginal genealogy has highest posterior probability under a Brownian motion dispersal process. In this sense, the genome-wide average reconstruction produced by our method can be viewed as a weighted average of posterior modes. Deraje et al. (2024) recently extended the model of branching Brownian motion to work with the full ARG rather than a sample of local gene trees.

---

**Figure S1.**
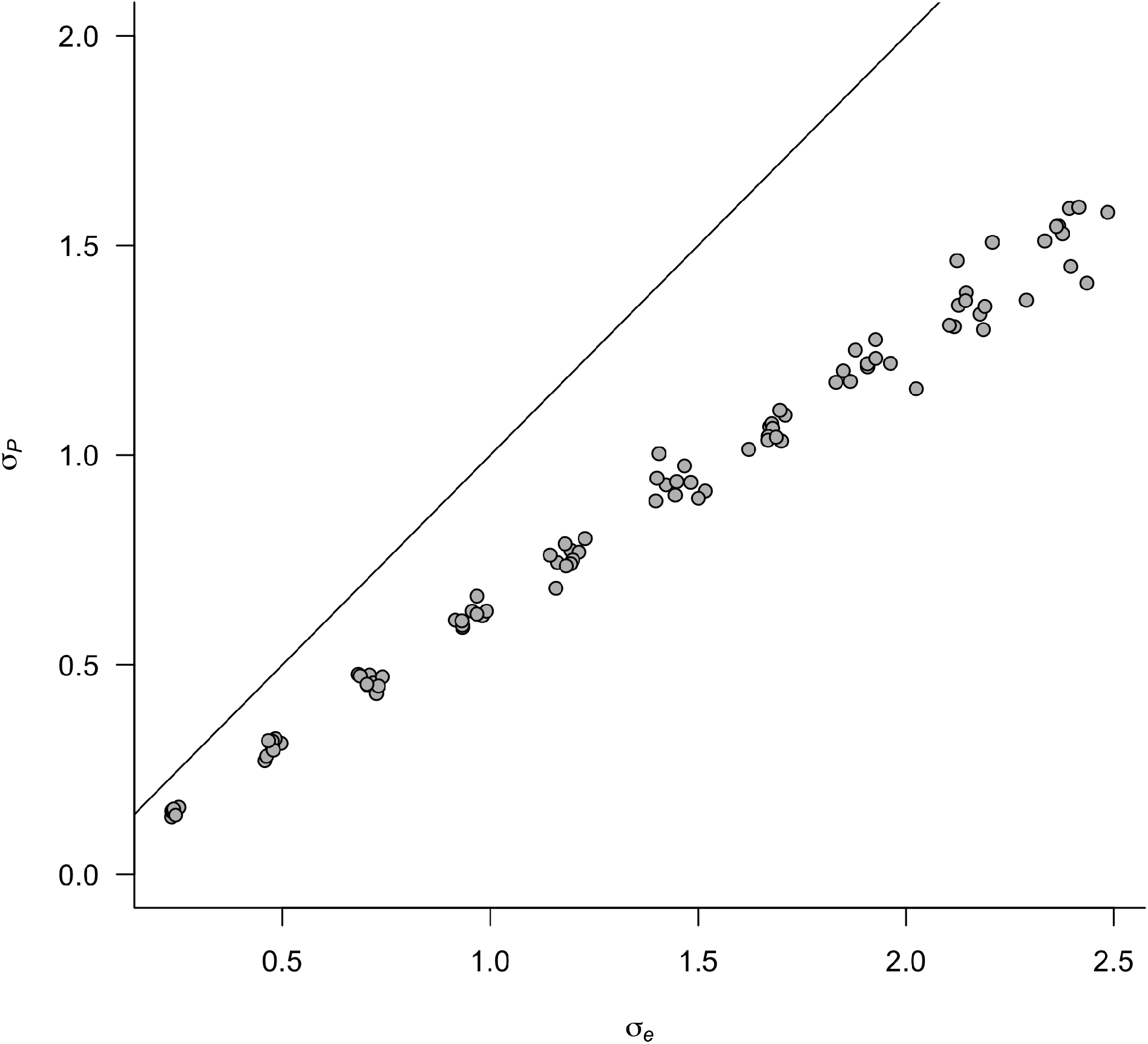
Migration rate estimates for a Gaussian dispersal kernel. Each point represents a single simulation generated under Gaussian dispersal with effective migration rate given on the x-axis and a parsimonious genome-wide estimate of that rate on the y-axis. The inset line shows a 1:1 relationship.

**Figure S2.**
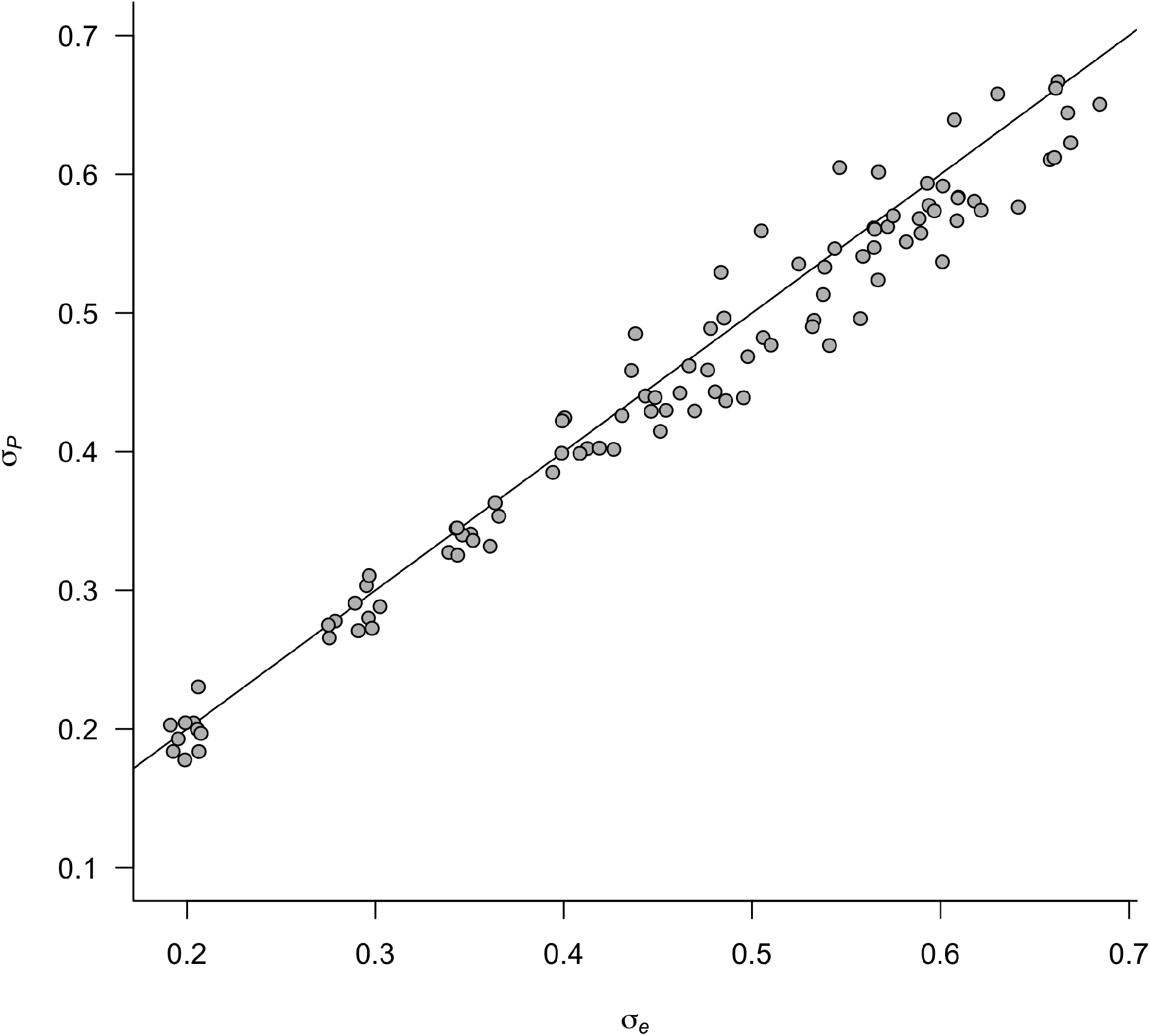
Migration rate estimates for a Laplace dispersal kernel. Each point represents a single simulation generated under Laplace dispersal with effective migration rate given on the x-axis and a parsimonious genome-wide estimate of that rate on the y-axis. The inset line shows a 1:1 relationship.

**Figure S3.**
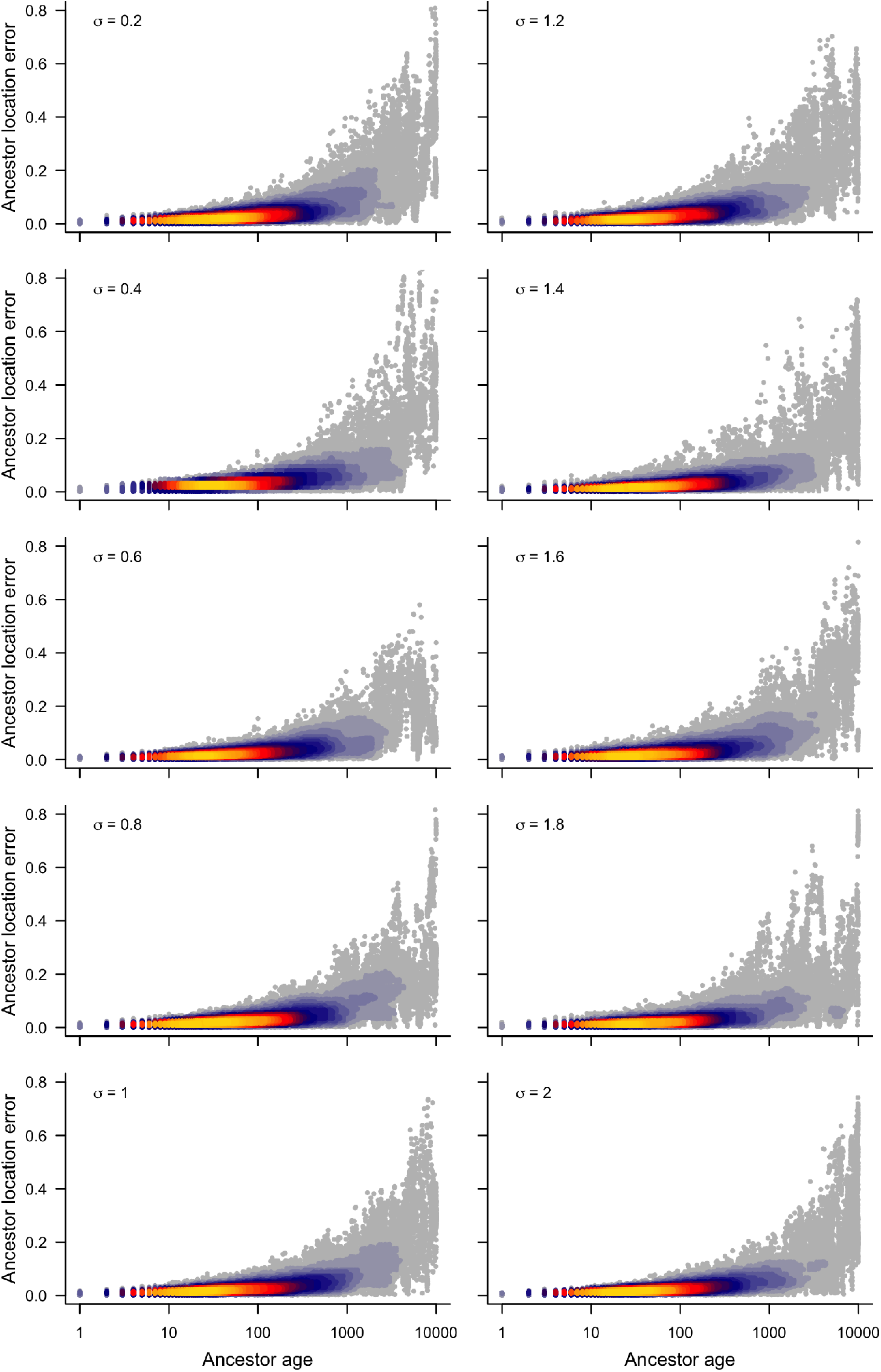
Ancestor location error for a Gaussian dispersal kernel. Each point represents a single genetic ancestor. Ancestor location error is measured as the distance between the estimated and the true location divided by the greatest distance separating any pair of samples. Warm colors signify a greater density of points.

**Figure S4.**
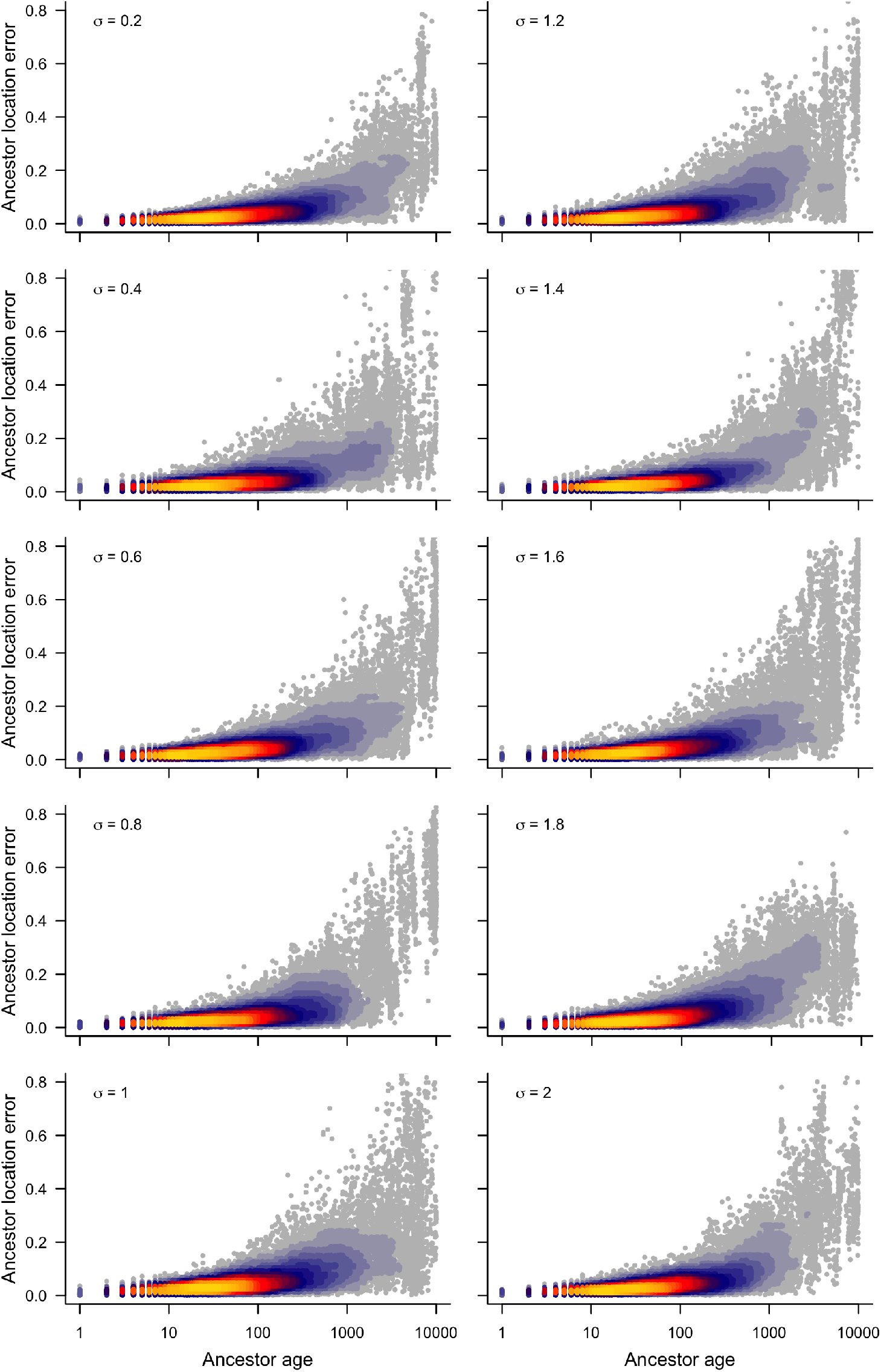
Ancestor location error for a Laplace dispersal kernel. Each point represents a single genetic ancestor. Ancestor location error is measured as the distance between the estimated and the true location divided by the greatest distance separating any pair of samples. Warm colors signify a greater density of points.

